# Directed manipulation of membrane proteins by fluorescent magnetic nanoparticles

**DOI:** 10.1101/2020.03.30.016477

**Authors:** Jia Hui Li, Paula Santos-Otte, Braedyn Au, Jakob Rentsch, Stephan Block, Helge Ewers

**Author notes:** **Corresponding author**: Helge Ewers, Institut für Biochemie, Freie Universität Berlin, Thielallee 63, 14195 Berlin, Germany.

## Abstract

The plasma membrane is the interface through which cells interact with their environment. Membrane proteins are embedded in the lipid bilayer of the plasma membrane and their function in this context is often linked to their specific location and dynamics within the membrane. However, few methods are available for nanoscale manipulation of membrane protein location at the single molecule level. Here, we report the use of fluorescent magnetic nanoparticles (FMNPs) to track membrane molecules and to manipulate their movement. FMNPs allow single-particle tracking (SPT) at 10 nm spatial and 5 ms temporal resolution, and using a magnetic needle, we pull membrane components laterally through the membrane with femtonewton-range forces. In this way, we successfully dragged lipid-anchored and transmembrane proteins over the surface of living cells. Doing so, we detected submembrane barriers and in combination with super-resolution microscopy could localize these barriers to the actin cytoskeleton. We present here a versatile approach to probe membrane processes in live cells via the magnetic control of membrane protein motion.

## Introduction

Membrane proteins are heterogeneously distributed in the plasma membrane of cells despite its fluid and continuous nature^1,2^. Local accumulation or separation of specific membrane molecules is crucial for efficient execution and regulation of fundamental biological processes such as (synaptic) signaling^3^, cellular secretion^4^ and internalization of nutrients^5^, cellular movement^6^ and tissue formation^7^. Imaging techniques such as super-resolution imaging, SPT, and fluorescence correlation spectroscopy have been instrumental to describe the local membrane composition and dynamics^8^. To perturb the functional localization and dynamics of a given membrane molecule, genetic and pharmacological means are usually chosen. However, these approaches lead to global effects and are very limited in temporal resolution. Optogenetics and other photoactivation methods enable better temporal control but nevertheless lack spatial information at the single molecule level. To directly move membrane molecules, optical tweezers^9,10^ have been used, but this method requires sophisticated equipment and mostly allows the control of only one particle at a time. Here, we sought to use magnetic nanoparticles as a straightforward alternative. Magnetic nanoparticles are widely used in biomedicine for imaging and therapy^11,12^. In biophysics, magnetic tweezers with micrometer-sized magnetic beads are established tools for force spectroscopy^13^. In the last decade, advances in the use of biofunctionalized magnetic particles to remotely control cellular processes have opened a new field termed “magnetogenetics”^14^. Micro- to nanometer-sized magnetic particles coated with active biomolecules are introduced to the specimen to actuate mechanical, thermal, or biochemical signals. At the plasma membrane, mechano-, heat-, or clustering-sensitive receptors and ion channels can be activated using magnetic particles^15–17^. Usually, either large clusters or a high concentration of particles are used for ensemble measurements. In contrast, we here use particles sized below the diffraction limit of light in a low-density regime to retrieve information on single molecules with high spatiotemporal resolution. The magnetic component of the particles then allows concomitant manipulation by applying an external magnetic field. Our method provides spatial control over membrane protein motion while determining its localization with nanometer precision. This allows for the exact correlation of membrane protein location with cellular events and structures.

## Results

First, we aimed to determine whether FMNPs were compatible with SPT and magnetic manipulation in the focal plane of a fluorescence microscope in aqueous conditions. To do so, we made use of particles consisting of a 100 nm diameter ferromagnetic core and a polymer shell conjugated with a fluorescent dye and streptavidin. We found these particles to be monodisperse in aqueous buffer and brightly fluorescent. The FMNPs bound readily to supported lipid bilayers (SLB) containing 0.001 mol% biotinylated lipids and were detectable as single, diffraction-limited spots (Fig. 1a). When we tracked the particles over time at a framerate of 50 Hz in TIRF illumination, we found that they exhibited random motion in the plane of the membrane with a mean/median diffusion coefficient *D* of 0.13/0.15 μm^2^/s (931 tracks; Fig. 1b, left; Supplementary Movie 1). When we then approached a magnetized needle^18^ to the field of view using a micromanipulator, the particles started to move towards the tip of the needle (Fig. 1b, middle; Supplementary Movie 1). Upon removal of the needle, the particles returned to random motion (Fig. 1b, right; Supplementary Movie 1). We concluded that we could pull lipids in the plane of the membrane into the direction of the needle via FMNPs and that we could do so in a reversible and temporally controlled manner.

**Figure 1:**
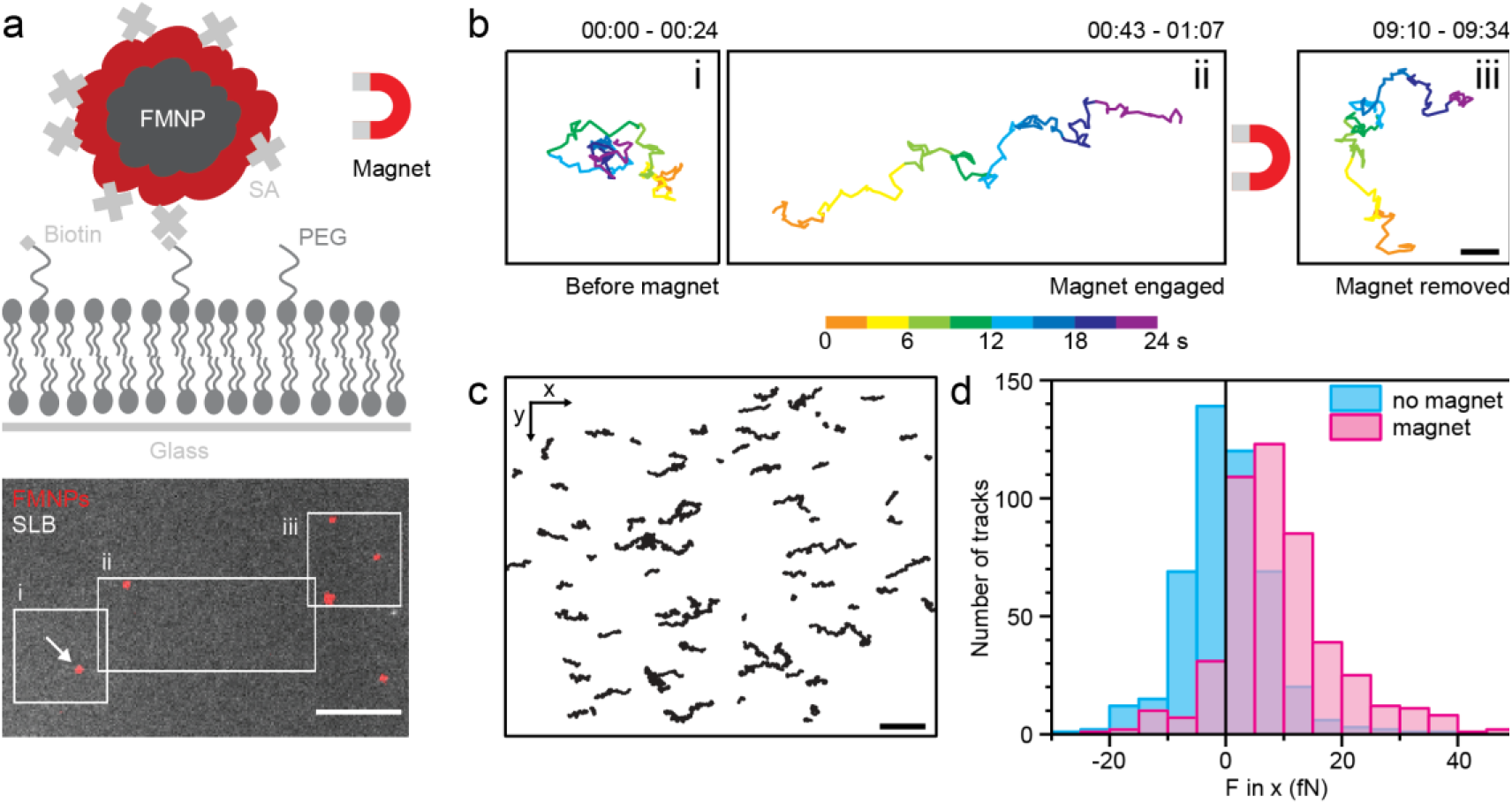
Manipulation of lipids by fluorescent magnetic nanoparticles (FMNPs) on a supported lipid bilayer (SLB) and force calibration. a) Top: Experimental setup for the directed manipulation of a Streptavidin (SA)-functionalized FMNP bound to biotinylated lipid (DSPE-PEG(2k)-biotin) in a SLB. Graphic is not to scale. Bottom: Overlay of the first frame of a fluorescence time-lapse video of FMNPs (red) bound to a SLB doped with carboxyfluorescein-conjugated lipid (DOPE-CF). Scale bar is 10 μm. b) Trajectory of the FMNP marked with an arrow in a) before (i), during magnetic manipulation (ii) and afterwards (iii) colored by time. Scale bar is 2 μm. c) Trajectories of FMNPs tracked on a SLB in presence of a magnetic needle to the right of the plot. d) Histogram of forces per trajectory exerted on FMNPs in presence (magenta, 469 tracks) and absence (blue, 460 tracks) of the magnetic needle.

The particle movement under magnetic attraction results from the directed motion towards the magnetic needle and the inherent Brownian motion. To estimate the forces that are exerted on the FMNP-bound phospholipid by the magnetic needle, we separated the components of the trajectory in-direction-of (directed component) and perpendicular-to (Brownian component) the needle tip. Using equation (1)^19^, we directly calculate the force from each trajectory of hundreds of particles bound to lipids in SLBs (Fig. 1c).

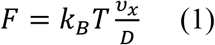

The average velocity *v*_x_ of any FMNP can be extracted from its linear movement in the direction of the magnetic field (x-axis) and the average FMNP diffusion coefficient *D* from the fluctuations in the direction perpendicular to the magnetic field (y-axis). The calculated forces in x-direction are 9 ± 11 fN (mean ± standard deviation, 469 tracks, Fig. 1d) when the magnetic field is applied and are undetectable without the magnet (0 ± 8 fN, 460 tracks).

Notably, both the directed and random movement components are apparent in the trajectories (Fig. 1b), i.e. the displacement per frame due to the applied force is comparable to the one due to Brownian motion. The energy which is spent to drag the molecule through the lipid bilayer against friction is therefore on the order of the thermal energy *k*_*B*_*T*. This is desirable when applied to membrane proteins on living cells as forces in the femtonewton range will not abrogate most naturally occurring interactions of membrane proteins^20^. Instead, the magnet-attracted membrane protein can probe its path for functional interactions in the membrane environment. We concluded that FMNPs work as a straightforward probe to track membrane-bound molecules and to move them in a controlled, directed manner.

To estimate the localization precision we achieved in fluorescence microscopy, we imaged immobile FMNPs on the SLB and localized them over thousands of subsequent frames. Because of the high fluorescence brightness, the position center could be determined with high precision by sub-pixel localization. We found that FMNPs could be localized with 10 nm precision (Fig. 2a), and this was possible at framerates down to 5 ms and up to 10,000 frames. To demonstrate this also for moving particles, we took advantage of structural inhomogeneities of the SLBs that were created by scratching the cover-glass resulting in a gap in the continuous lipid bilayer. When we pulled FMNPs attached to biotinylated lipids in the SLB against such defects, we found that they appeared to slide along the defect boundaries (Fig. 2b). Since the FMNPs can be localized with nanometer localization precision, they map the boundaries of the SLB significantly more accurate than the diffraction-limited image of the SLB (image resolution > 200 nm, Fig. 2c).

**Figure 2:**
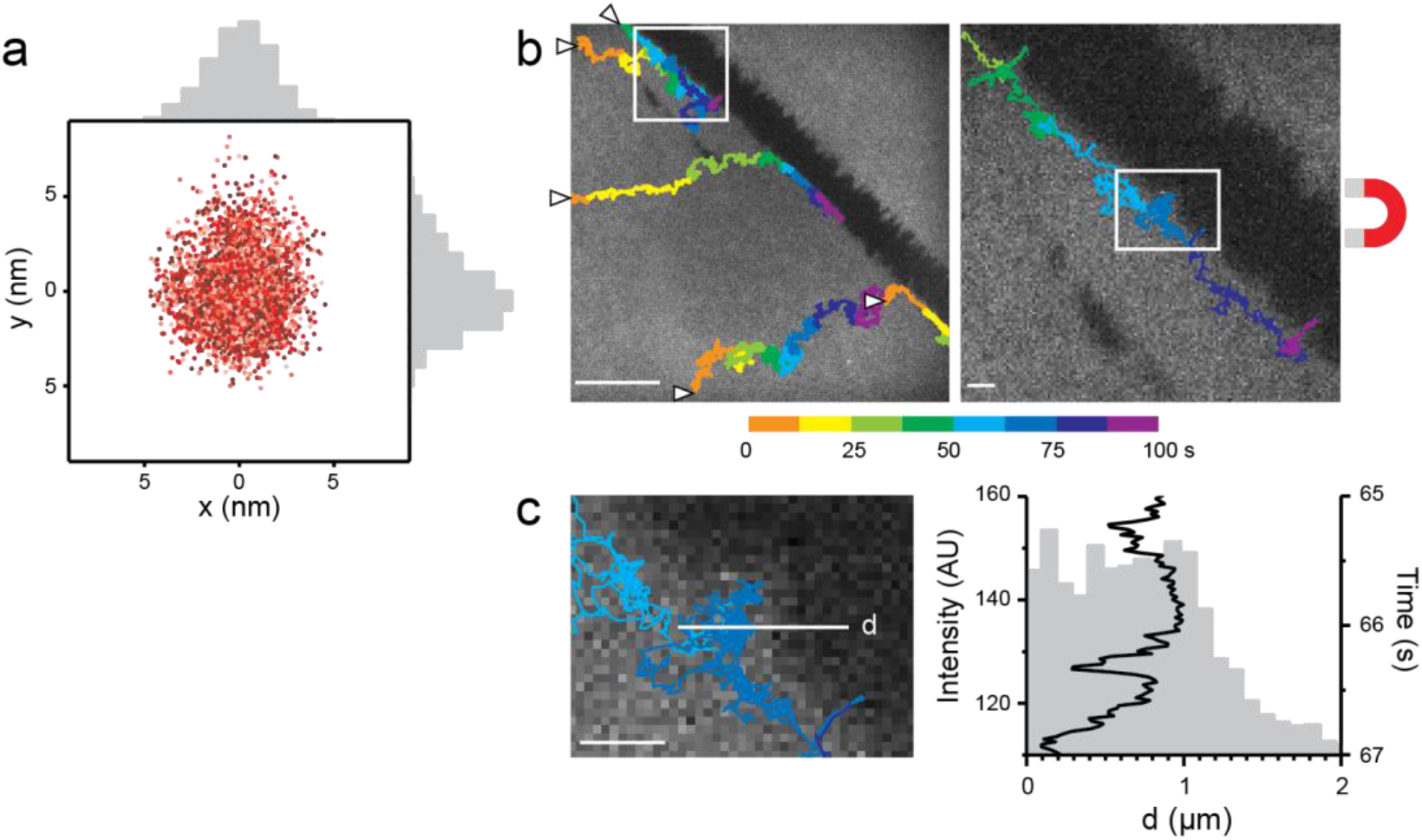
Accuracy of FMNP localization and tracking. **a)** Plot of subsequent 2D localizations of a single immobile FMNP on a SLB by subpixel fitting in each frame (total 3000 localizations, imaged at 50 Hz), colored by time (light to dark red). **b)** Trajectories of SLB-bound FMNPs under magnetic pull (beginning marked with arrowheads) overlaid onto the fluorescence image of the SLB (grey) with defects (black areas). Scale bar is 10 μm. **c)** Fluorescence micrograph of inset in b) and plot of FMNP localization in direction of the white line (d) over time above plot of SLB fluorescence intensity along (d). Scale bar is 1 μm.

We next aimed to test whether our system could be applied to living cells. To do so, we coupled FMNPs to commonly studied plasma membrane proteins transiently expressed in living mammalian cells. We expressed a probe for the outer leaflet of the plasma membrane, glycosylphosphatidylinositol-anchored green fluorescent protein (GPI-GFP), and two transmembrane probes, the yellow fluorescent protein-tagged single-spanning transmembrane domain of the low density lipoprotein-receptor (L-YFP-GT46) and the GFP-tagged full-length transferrin-receptor (TfR-GFP). To deliver the FMNPs to the target probe, we used biotinylated anti-GFP nanobodies, which bind to GFP and derived variants such as YFP (Fig. 3a). The nanobody-conjugated FMNPs bound specifically on the dorsal membrane of transfected cells (Supplementary Figure 1) and single, mobile FMNPs could readily be tracked. When we approached the sample with the magnetic needle, the outer-leaflet probe as well as the transmembrane proteins could be moved over several micrometers in the plane of the plasma membrane towards the needle (Fig. 3b). We concluded that we could track and manipulate proteins with different membrane anchoring moieties on living cells using FMNPs.

**Figure 3:**
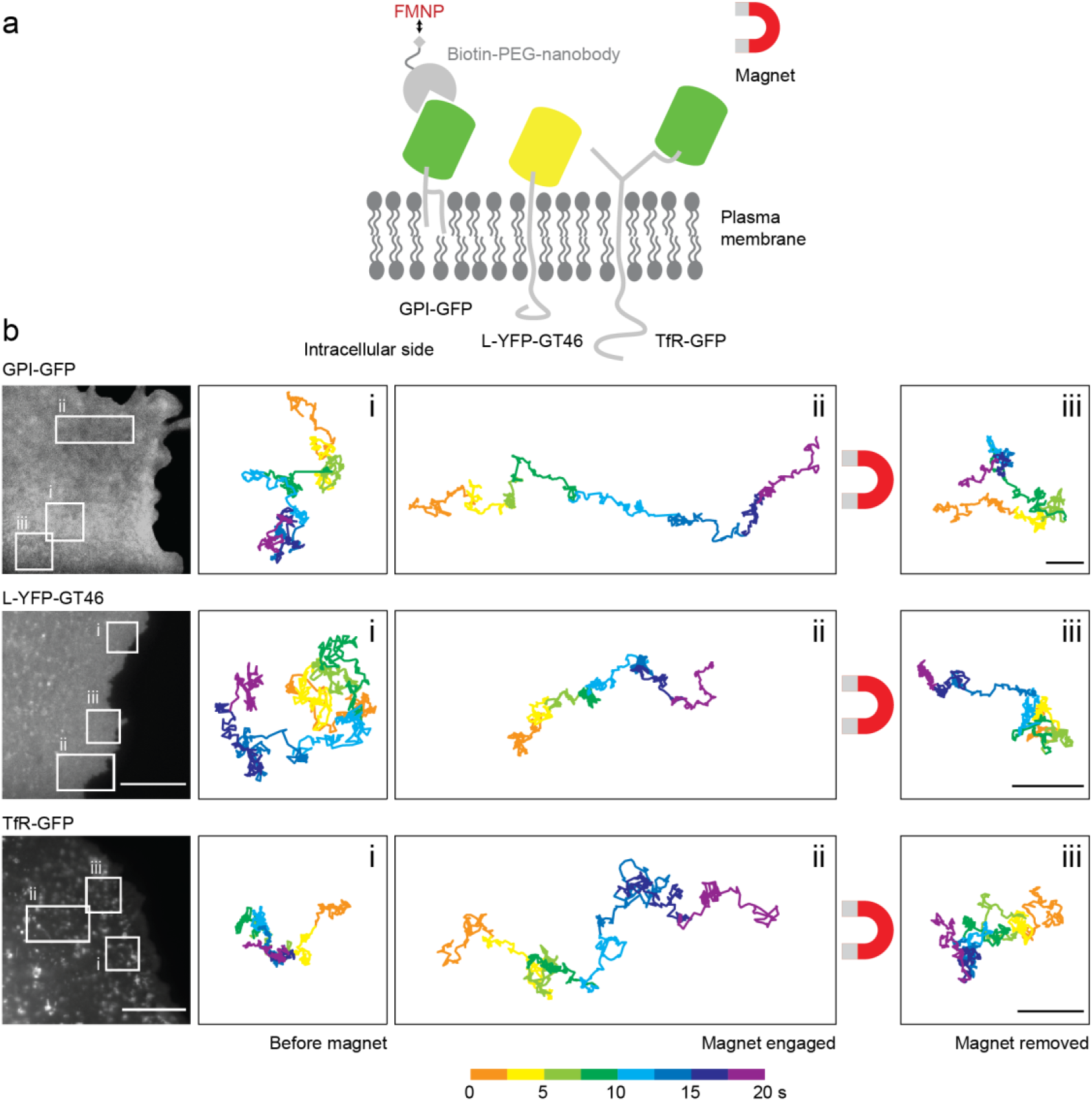
Manipulation of membrane proteins by FMNPs on the surface of living cells. **a)** Scheme of the manipulation of the outer leaflet lipid-anchored (GPI-GFP) and transmembrane protein probes (L-YFP-GT46 and TfR-GFP) in living cells. The FMNPs are targeted to the fluorescent protein moiety of membrane protein probes via biotinylated anti-GFP nanobodies and then pulled across the plasma membrane by a magnetic needle. Graphic is not to scale. **b)** From top to bottom: Experiments of membrane protein probes pulled across the plasma membrane of living CV-1 cells expressing GPI-GFP, L-YFP-GT46, or TfR-GFP. Left: Fluorescence micrographs of single cells expressing the respective constructs with areas in which protein-bound FMNPs were imaged before (i), during (ii) and after (iii) magnet engagement. Scale bars are 10 μm. Right: Trajectories of exemplary FMNPs bound to the respective membrane probes in the white-framed areas on the cell under the magnetic conditions (i-iii). Trajectories are color coded for time. Scale bars are 1 μm.

We then aimed to correlate the motion of membrane proteins with the location of cellular structures. In living cells, the cortical cytoskeleton affects the movement of membrane lipids and proteins^21–27^. Cytoskeletal actin-protein polymers form a dense meshwork underneath the plasma membrane^28^. To ask whether we could detect a physical barrier to membrane protein motion formed by actin filaments (F-actin), we pulled GPI-GFP molecules over the surface of CV-1 cells, fixed them and subsequently performed biplane 3D (*d*)STORM super-resolution microscopy of F-actin via phalloidin-AF647. This allowed us to specifically visualize cortical F-actin at the dorsal membrane (Supplementary Figure 2). We then used fiducial markers present both in the SPT and (*d*)STORM experiments to overlay the SPT trajectory with the (*d*)STORM super-resolution image of the cortical F-actin. Doing so, we could detect that a GPI-GFP molecule, which was dragged for > 2 μm across the cell membrane, slowed down and came to a halt (Fig. 4a, Supplementary Movie 2). The particle did not move directly towards the needle but seemed to avoid sites of dense F-actin (Fig. 4b). It then slowed down as it approached a thick, bundled actin filament and slid along this filament in direction of the needle. This movement pattern suggests that the GPI-GFP either experienced a physical barrier, directly imposed by the actin filament or indirectly by actin-associated transmembrane proteins or extracellular matrix, or a specific interaction with a cellular structure such as a clathrin-coated pit. On a broader scope, this experiment demonstrates that sites of interaction can be detected immediately and dynamically by introducing a directed movement component to the diffusional motion. While classical SPT experiments also allow the exploration of the local membrane structure, due to the random nature of molecular diffusion, robust conclusions can only be drawn given high statistical sampling in time and space. In contrast, our approach makes it possible to directly probe the interaction of a membrane molecule with specific cellular structures and to obtain meaningful results for each individual particle.

**Figure 4:**
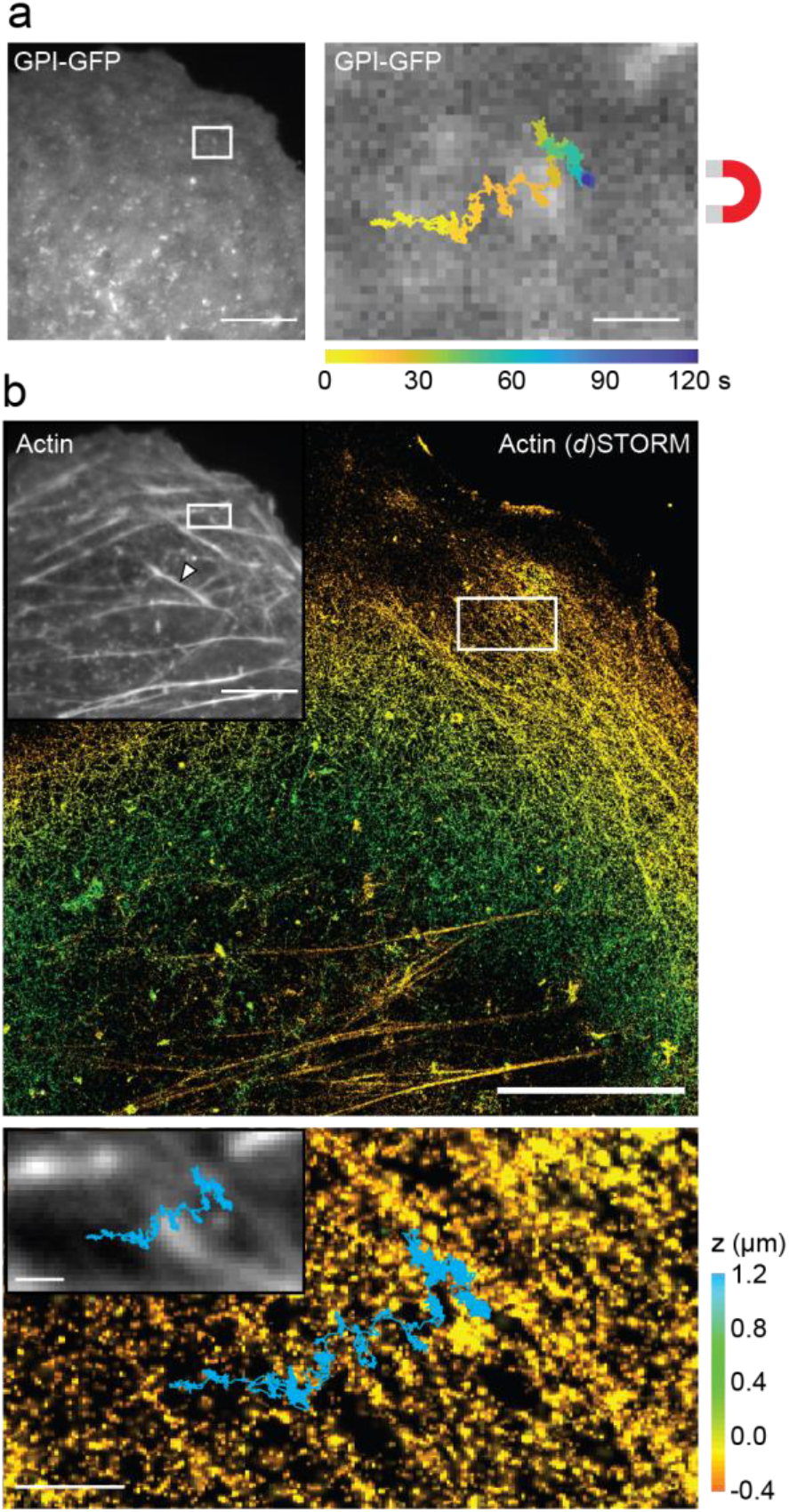
Correlation of FMNP-steered protein movement on the plasma membrane and the underlining F-actin cytoskeleton. **a)** Live fluorescence image of the membranous GPI-GFP signal in a CV-1 cell. Scale bar is 10 μm. Right: Trajectory of a FMNP bound to GPI-GFP and pulled by the magnetic needle overlaid onto the GPI-GFP signal. Scale bar is 1 μm. **b)** Top: Phalloidin-AF647 stained F-actin in the same cell as in a) after fixation, imaged by conventional epifluorescence microscopy (inset) and 3D (*d*)STORM super-resolution microscopy (color scale by depth z, bottom 0.6 μm z-slice not included). Arrowhead in the epifluorescence inset indicates a stress fiber which is excluded in the super-resolution z-slice. Scale bar is 10 μm. Bottom: Trajectory of FMNP (cyan) overlaid onto the epifluorescence (inset) and (*d*)STORM image after image registration via fiducials. Scale bar is 1 μm.

## Discussion

By using FMNPs, we have here combined the non-invasive perturbation capabilities of magnetic fields with high-precision SPT of a molecule of interest. Compared to optical tweezers, our approach is straightforward, low-cost, and allows to move several FMNPs at the same time. Only at very high densities it must be considered that particles may attract each other under magnetization and aggregate. Due to our lateral magnetic tweezer setup, the pulling force can be applied in the focal plane of conventionally used inverted microscopes^29^. Previous ensemble applications of magnetic nanoparticles have already demonstrated the potential of local control over cellular processes^15,18^. Our present approach now combines such magnetic control of movement with SPT to enable experiments at high resolution in time and space. This approach could be extended to other, non-fluorescent SPT techniques such as interferometric scattering microscopy (iSCAT)^30^ for even faster measurements. Furthermore, we demonstrate the compatibility with other imaging techniques by performing post-hoc super-resolution imaging of the cytoskeleton. In principle, the magnetic manipulation could be performed concomitantly with other live super-resolution imaging methods such as photoactivated localization (PALM) or live-cell stimulated emission depletion (STED) to capture the dynamic remodeling of the cellular environment. The combination of these techniques enables the interrogation of membrane components and their local environment at spatiotemporal scales that match their activity. In this context, exerting only small forces on the molecule of interest is crucial to maintain the potential for productive intermolecular interactions. In the future, this method could be applied to study receptor signaling at the plasma membrane. One could extract a specific component of a signaling hub and monitor the downstream effects in real time. Receptor dimerization for example is an important but hardly tractable problem in signaling. By pulling via an FMNP on one receptor subunit and simultaneously imaging the movement of the other subunit, quantitative data on the dimerized fraction of receptors in the membrane should be instantly accessible. Another field of application could be the investigation of polarized cells in which different membrane domains have a distinct functional organization and composition. A molecule of interest could be moved to an ectopic location on the membrane to study its context-dependent activity. Combining SPT and magnetic manipulation expands our current toolbox of methods to observe and to perturb the location of molecules in the plasma membrane. Our work thus opens the doors to easily accessible experiments aimed at understanding the interaction of membrane molecules with specific membrane-apposed cellular structures.

## Material and methods

### Nanoparticle characterization and magnetic tweezer setup

100 nm BNF-Starch-redF particles with Streptavidin surface (Micromod) are composed of a magnetite core and a starch shell crosslinked with the red fluorescent dye Dy555. They are thermally blocked at room temperature. Dynamic light scattering measurements with a 633 nm laser (Zetasizer Nano, Malvern) of the particles in PBS confirmed that they are monodisperse with a polydispersity index of 0.11 (< 0.2 is considered monodisperse) and a hydrodynamic radius of 159 ± 2 nm (Z-Average size). The magnetic needle^18,31^ is made of a spring steel wire of 0.1 mm diameter (Fohrmann-Werkzeuge), which was pulled in a Bunsen flame to create a tip. The needle was then glued to a 6×4×2 mm Neodyn magnet (QM-06×04×02-N, Magnets4you) and attached to a motorized micromanipulator (PatchMan, Eppendorf). The magnetic needle was lowered until it touched the bottom of a dummy sample dish and raised again so that it was placed above the bottom of the sample with the very tip in focus around 100 μm above the focus of the glass surface.

### SLB preparation and particle binding

Small unilamellar vesicles were prepared by sonication and used to form SLBs by the vesicle drop method. DOPC (1,2-dioleoyl-sn-glycero-3-phosphocholine), DSPE-PEG(2k) (1,2-distearoyl-sn-glycero-3 phosphoethanolamine-N-[carboxy(polyethylene glycol)-2000]), DSPE-PEG(2k)biotin (1,2-distearoyl-sn-glycero-3-phosphoethanolamine-N-[biotinyl(polyethylene glycol)-2000]) and DOPE-CF (1,2-dioleoyl-sn-glycero-3-phosphoethanolamine-N-carboxyfluorescein) were purchased from Avanti Polar Lipids. DOPC doped with 0.2-2 mol% PE-CF and 1-2 mol% of either DSPE-PEG(2k) or DSPE-PEG(2k)-biotin were dissolved in 1:1 chloroform:methanol and dried under vacuum overnight. The lipids films were hydrated in SLB buffer (2 mM CaCl_2_, 200 mM NaCl and 10 mM HEPES, pH 6.8; all Sigma Aldrich), sonicated for 50 s and centrifuged for 30 min at top speed. The supernatants were stored at 4°C for up to 3 weeks until use. Round 25 mm #1.5 coverslips (VWR) or 35 mm diameter glass-bottom dishes (Zell-Kontakt) were plasma cleaned for 5 min prior to SLB formation. Solutions of DSPE-PEG(2k)- and DSPE-PEG(2k)-biotin-containing vesicles were mixed in appropriate concentration ratios to achieve a final concentration of 0.001-0.005 mol% DSPE-PEG(2k)-biotin in the solution and dropped directly onto the glass or into a 4-well silicon mold (Ibidi) on the glass. The bilayer was washed ten times with SLB buffer and evaluated on the spinning disc confocal microscope by fluorescence recovery after photobleaching (FRAP, data not shown). SLBs were incubated with 1×10^9^/ml FMNPs in SLB buffer with 1% BSA on the microscope until tens to hundreds of particles per field of view had bound to the SLB, and then washed 5-10 times with SLB buffer before imaging.

### Cell culture

Wildtype *Cercopithecus aethiops* kidney fibroblasts (CV-1) were grown in DMEM without phenol red, supplemented with 10% fetal bovine serum (FBS) and GlutaMAX (all Thermo Fisher Scientific) at 37°C in a CO_2_-controlled humidified incubator. Stably transfected GPI-GFP CV-1 cells were maintained with 50 μg/ml Geniticin (Thermo Fisher Scientific) supplementing the culture medium. For transient transfection, CV-1 cells at 70-100% confluency were electroporated with the Neon Transfection System according to the supplier’s protocol for COS-7 cells. Cells were plated on glass-bottom dishes and imaged in imaging medium (Fluorobrite DMEM, 10% FBS, GlutaMAX, 10-30 mM HEPES; Thermo Fisher Scientific) 16-48 h after transfection.

### Particle functionalization and live cell experiments

For live cell experiments targeting GFP or YFP, LaG-16 anti-GFP nanobodies^32^ were recombinantly produced in house and labeled with α-Biotin-ω-(succinimidyl propionate) 24(ethylene glycol) (Iris Biotech) as previously described^33,34^. The succinimidyl ester was added to the nanobody at 2-fold molar excess for the labeling reaction. After purification via Zeba Spin Desalting Columns (Thermo Fisher Scientific), the labeling efficiency was estimated to be 0.06 dye/protein by labeling with Alexa Fluor 647 carboxylic acid, succinimidyl ester in parallel. The FMNPs were mixed with the biotinylated nanobody in a 10:1 to 1:2 molar ratio, magnetically separated for 1 hour to remove the majority of unlabeled nanobodies, and then resuspended in imaging medium. Living cells were incubated with this suspension at a final concentration of 10^10^-10^11^ particles/ml in imaging medium for 15 min, washed 3-5 times with PBS or medium, and imaged in imaging medium.

### Microscopy

SLBs were imaged with TIRF illumination. Cells were imaged in epifluorescence illumination because the FMNPs bind to the apical cell membrane. Typically for SPT, a single image series of 1,000-10,000 frames was collected with a 5-50 ms acquisition time (20-200 Hz, in total 20 s to 3:20 min) at a laser-power density of 8-200 W cm^−2^ using the 561 nm laser. All experiments were carried out at room temperature to reduce endocytic activity. For the force calibration, longer frame times were recorded (100 ms exposure time with 1-7 Hz frame rate). For correlative imaging of the cytoskeleton, 100-nm red-fluorescent (580/605) beads (Thermo Fisher Scientific) were added as fiduciary markers and cells were fixed immediately after live-cell imaging and stained with phalloidin-Alexa Fluor 647^28^. (*d*)STORM imaging was performed as described previously^35^ at a laser-power density of 4.4 kW cm^−2^ using the 640 nm laser, acquiring 20,000-40,000 frames with 20 ms acquisition time. SPT and (*d*)STORM imaging were performed on a Vutara 352 super-resolution microscope (Bruker) equipped with a 60×/NA 1.49 oil immersion TIRF objective (Olympus) and an ORCA Flash 4.0 V2 CMOS (Hamamatsu). The force calibration experiments were performed on an inverted Ti-E TIRF microscope (Nikon) that was equipped with a high-pressure mercury lamp, an Apo TIRF 60×/NA 1.49 oil objective, and an Andor Zyla 4.2 sCMOS camera (Andor Technology). FRAP was performed on an inverted IX71 microscope (Olympus) equipped with a CSU-X1 spinning disk (Yokogawa) and an iLas2 FRAP system (Gataca Systems). A 60×/NA 1.42 oil objective (Olympus) was used and time-lapse image sequences were taken with an ORCA Flash 4.0LT CMOS camera (Hamamatsu).

### Data analysis

Analysis of (*d*)STORM imaging was done in the SRX software (Bruker). The localizations were rendered with a Gaussian distribution by radial precision (Euclidean norm of the precision of the x and y axes calculated as the Cramér-Rao lower bound from the Fisher information of the point spread function). All other images were analyzed with Fiji^36^. SPT analysis was performed with the MOSAIC Particle Tracker^37^ or with Trackmate^38^. Immobile particles (*D* < 0.005 μm^2^/s) and trajectories shorter than 50 frames were discarded. Image registration via fiducials was performed by *nonreflective similarity* transformation in Matlab (MathWorks) using the built-in function *fitgeotrans*. Forces were calculated with a Matlab script adapted from Block et al.^19^. For the force calculation, trajectories slower than 0.01 μm^2^/s and shorter than 50 frames were discarded. Trajectories and other data were analyzed and plotted with OriginPro (OriginLab).

## Acknowledgements

The authors thank Prof. Blankenstein and Prof. Willemsky for lending the micromanipulator and the engineering workshop of the Institute of Chemistry of the Freie Universität Berlin for the custom fabrication of adaptors for the micromanipulator. The GPI-GFP and TfR-GFP plasmids were a kind gift from the Helenius laboratory, the L-YFP-GT46 plasmid was a kind gift of Patrick Keller. We thank Areeya Chawandee, Raluca Groza, and Dr. Vladimira Petrakova for testing reagents or performing initial experiments. The work was funded by Deutsche Forschungsgemeinschaft (DFG) through SFB 958, SFB 756 and TRR186. The authors acknowledge technical support from Andrea Senge, Claire Schlack, and Reinmar Undeutsch.

## Author contributions

H.E. designed research; J.H.L. and P.S.O. performed research; B.A., J.R. and S.B. contributed new reagents or analytic tools; J.H.L analyzed data; J.H.L and H.E. wrote the manuscript with input from all authors.

**The authors declare no conflict of interest**.

