## Supplementary figures and video legends for "Directed manipulation of membrane proteins by fluorescent magnetic nanoparticles"

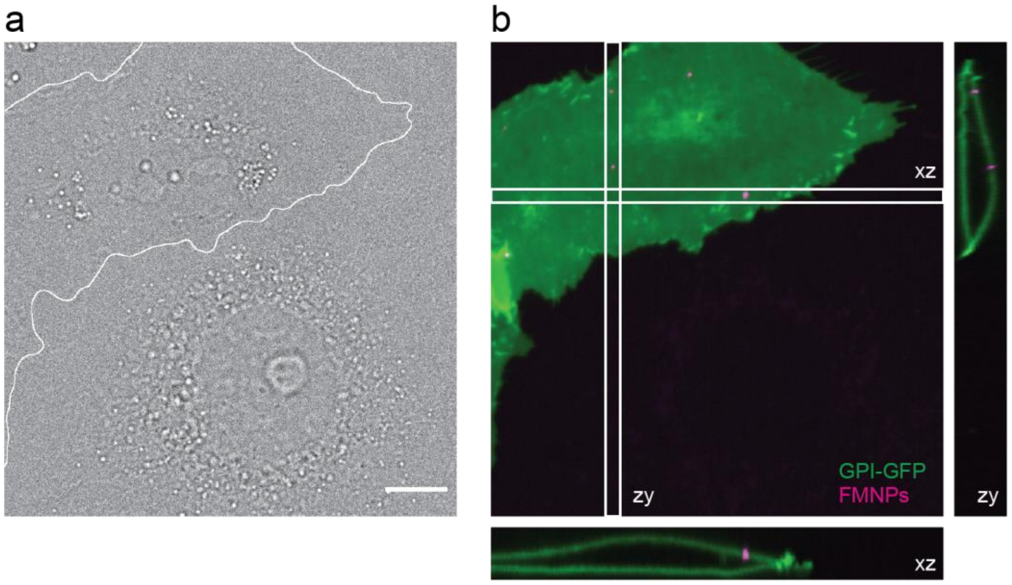


**Supplementary Figure 1: FMNPs are specifically targeted to GFP-expressing cells by GFP nanobodies. a)** Transmission light micrograph of a GPI-GFP expressing (white outlined) and a wild-type CV-1 cell. Scale bar is 10 µm. **b)** Average projection of a fluorescence z-stack of a) shows that the nanobody-coated FMNPs (magenta) have bound specifically to GPI-GFP (green). Orthogonal views (zy and xz) demonstrate the binding to the dorsal side of the cell.

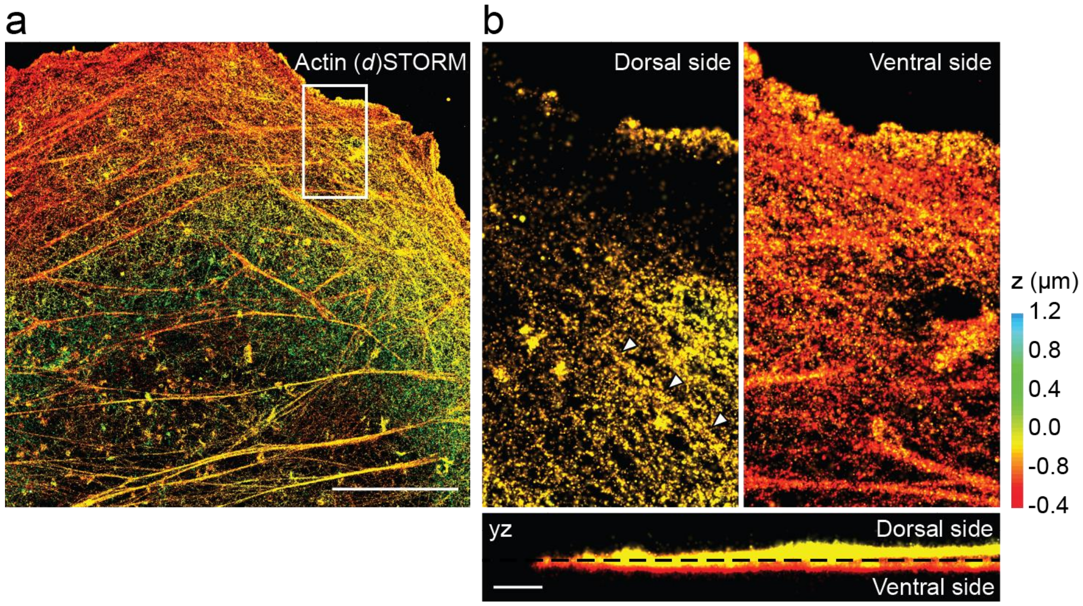


**Supplementary Figure 2: The actin filament shown in Fig. 4 is located at the dorsal side of the plasma membrane. a)** Full z-range reconstructed (*d*)STORM image of F-actin of the same cell as in Fig.4. Scale bar is 10 µm. **b**) Close-up view of selected z-slices of the dorsal (left) and ventral (right) cortical F-actin, as indicated in the yz view below. Scale bar is 1 µm.

**Supplementary Movie 1: Reversible magnetic manipulation of a FMNP-bound lipid in a supported lipid bilayer.** The same particle as in Fig. 1a (white arrow) bound to DSPE-PEG(2k)-biotin in a SLB was tracked before, during, and after magnetic manipulation. The magnetic tip was placed to the right side of the field of view. Single-particle trajectories are colored by time and overlaid onto the raw TIRF video. Playback speed is 5-fold of the real time speed.

**Supplementary Movie 2: A single FMNP bound to GPI-GFP on a live cell became immobile under magnetic manipulation.** The particle moved to the right side of the field of view where the magnetic tip was placed, but then slowed down and finally became immobile. The movement is correlated with the cortical F-actin cytoskeleton as shown in Fig. 4. Single-particle trajectory colored by time and overlaid onto the raw fluorescence time-lapse video. Playback speed is 10-fold of the real time speed.
